# Robustness of phylogenetic inference to model misspecification caused by pairwise epistasis

**DOI:** 10.1101/2020.11.17.387365

**Authors:** Andrew F. Magee, Sarah K. Hilton, William S. DeWitt

**Affiliations:** Department of Biology, University of Washington, Seattle, USA; Department of Genome Sciences, University of Washington, Seattle, USA; Department of Computational Biology Program, Fred Hutchinson Cancer Research Center, Seattle, WA

## Abstract

Likelihood-based phylogenetic inference posits a probabilistic model of character state change along branches of a phylogenetic tree. These models typically assume statistical independence of sites in the sequence alignment. This is a restrictive assumption that facilitates computational tractability, but ignores how epistasis, the effect of genetic background on mutational effects, influences the evolution of functional sequences. We consider the effect of using a misspecified site-independent model on the accuracy of Bayesian phylogenetic inference in the setting of pairwise-site epistasis. Previous work has shown that as alignment length increases, tree reconstruction accuracy also increases. Here, we present a simulation study demonstrating that accuracy increases with alignment size even if the additional sites are epistatically coupled. We introduce an alignment-based test statistic that is a diagnostic for pair-wise epistasis and can be used in posterior predictive checks.

## Introduction

Epistasis is the phenomenon where the effect of a mutation at one site in a sequence is dependent on the identity of another site or sites. This dependence is pervasive in datasets for phylogenetic inference and can manifest as interactions between different genes (Cohen et al., 2012; Schubert et al., 2019), between different sites in a protein (Dimmic et al., 2005; Rodrigue et al., 2009; Kryazhimskiy et al., 2011), or between different sites in RNA molecules (Shapiro et al., 2006; Nasrallah and Huelsenbeck, 2013; Meyer et al., 2019; Golden et al., 2020). Phylogenetic models of epistasis (and more broadly co-evolving sites) have focused on models of pairwise interactions within a single locus. While there has been work on models with larger alphabets (amino acids and context-sensitive mutation models) and high-order epistasis (Robinson et al., 2003; Rodrigue et al., 2009), these models are computationally burdensome, spurring the development of approximate computational approaches in order to fit them (Hwang and Green, 2004; Saunders and Green, 2007; Rodrigue et al., 2007; Laurin-Lemay et al., 2018). Phylogenetic models of pairwise epistasis include both general models of pairwise of interactions (Dib et al., 2014; Meyer et al., 2019) and explicit models of RNA evolution (Nasrallah and Huelsenbeck, 2013; Golden et al., 2020). In the case of RNA evolution, stem and loop secondary structures are conserved by paired substitutions at specific sites to maintain Watson-Crick pairing (A-T and G-C). Relaxing the independence assumption in this setting of pairwise interactions requires two new pieces of information: specifying the paired sites and a model of coupled character state change for pairs.

The issue of site pairings poses a practical difficulty in using an epistatic phylogenetic model for these pairings must be defined *a priori* or integrated out during inference. Defining pairings before analysis requires that such information is available in a database (*e.g.*, Wuyts et al., 2004), or that one can infer the RNA secondary structure from the sequence itself (*e.g.*, Lorenz et al., 2011). Approaches to integrate out the pairings are computationally costly. Common models that assume sites are independent and identically distributed (site-iid models) require computing the likelihood for each of *n* sites. To integrate out pairings, one could calculate the likelihood for all possible site pairings, as in Golden et al. (2020), but this requires 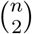 computations instead of *n*, greatly increasing run time. Meyer et al. (2019) employs reversible-jump MCMC (Green, 1995) to sample possible pairings, but the number of possible pairing schemes grows faster than *n*!, requiring longer runs to adequately explore the pairings.

Unmodeled pairwise interactions have been shown to decrease the accuracy of phylogenetic inference in simulations (Nasrallah et al., 2011). That is to say, phylogenies estimated from alignments simulated with epistatic interactions are farther from the true phylogeny than phylogenies estimated from alignments simulated without epistasis. Unmodeled epistasis could cause such differences in two ways. These differences could arise from bias caused by unmodeled epistasis (such bias is sometimes referred to systematic error, see for example Jeffroy et al. (2006)). In this scenario, inference with a model that ignores epistasis is expected to become more accurate if interacting sites are identified and omitted from the alignment. However, it is also possible that unmodeled epistasis *reduces the effective number of sites* in the alignment and increases the estimator variance without introducing bias. In this scenario, estimates are expected to worsen if the interacting sites are identified and omitted. As an example, consider two alignments, one of which has all sites simulated without epistasis, the other of which is the same as the first except that a proportion of sites are erroneously duplicated (copy-pasted into “new”” alignment columns). This is akin to the larger alignment being drawn from a particularly extreme model of pairwise epistasis. The larger alignment contains no new information, so the expected accuracy of the inferred phylogenies is the same for both alignments. However, compared to alignments of equal size, but where all sites were simulated without pairings, this larger alignment has reduced accuracy. It is effectively a shorter alignment. In this case, if one could remove one site from each pair inference should largely remain unchanged. In less extreme cases where sites are not perfect duplicates, removing half of the paired sites would remove some new information and thus decrease the accuracy of inference.

A number of methods have been proposed for detecting epistasis in multiple sequence alignments. This literature is largely based on mapping substitutions along a phylogeny and examining patterns of substitution. Shapiro et al. (2006) devise a test for epistasis based on multiple co-occurrences of substitutions along a phylogeny while Kryazhimskiy et al. (2011) use the order of substitutions through time to test for epistasis. However, these approaches, and others which map substitutions along a phylogeny, often ignore a potentially large source of uncertainty by conditioning on a single, estimated phylogney. Poon et al. (2007) propose a bootstrap procedure to account for phylogenetic uncertainty, and Dimmic et al. (2005) use a fully Bayesian approach to integrate out the phylogeny. A promising yet unexplored approach is to use posterior predictive checks of model performance as both the detection of epistasis and evaluation of model fit may be addressed simultaneously.

Given the pervasiveness of epistasis in real data and the difficulties involved in applying epistatic phylogenetic models to datasets, we seek to understand the cost of employing non-epistatic models to datasets with pairwise epistasis using simulations. In this paper, we ask two questions regarding the use of standard models, where sites are independent and identically distributed, in a misspecified setting where the alignment is generated from an evolutionary process with pairwise epistasis. First, can we detect the presence of unmodeled pairwise epistasis in datasets using posterior-predictive model checks? Second, what is the effect of including epistatically paired sites on the quality of trees inferred with a site-independent model?

To address these questions, we perform a simulation study using the pairwise epistatic model of RNA from Nasrallah and Huelsenbeck (2013). Briefly, this model aims to capture a common mutational process among all sites while implicitly accounting for the effect of selection against single mutations at predefined paired stem sites, which would break the secondary structure. We simulate alignments on a 3-dimensional parameter grid, defined by the number of sites in the alignment that are site-iid *n*_i_, the number of sites that are epistatically paired *n*_e_, and the strength of epistatic interactions *d* (the relative rate of secondary-structure-preserving double mutations to single mutations at paired sites). Our grid thus includes a range of alignment sizes *n* = *n*_i_ + *n*_e_ and a range of epistatic fractions *n*_e_/*n* for each size.

We assess several alignment test statistics—one previously described and several new ones designed to detect epistasis specifically—for their ability to detect epistasis directly from alignments using posterior predictive checks. This allows us to address our first question, can we *detect* pairwise epistasis? Next, we examine whether adding epistatic sites to an alignment makes inference better or worse. This allows us to address our second question, whether the inclusion of epistatic sites in an alignment improves or worsens phylogenetic estimates. We seek to quantify this second effect by estimating the relative worth, *r*, of an epistatic site in the alignment to a site drawn from a site-independent model. Using this concept of relative worth, we define two scenarios under our regime of model misspecification. In the best-case scenario (*r* > 0), the epistatic sites contribute useful information that improves phylogenetic inference while in the worse-case scenario (*r* < 0) the inclusion of epistatically paired sites makes inference worse. Finally, we combine the two questions and address whether pairwise epistasis is detectable when it is strong enough to significantly impact tree inference.

## Results

### Alignment-based test statistics are sensitive to pairwise interactions between sites

First, we use simulations to evaluate the sensitivity of alignment-based test statistics to pairwise interactions between sites. This sensitivity is necessary for the posterior predictive tests we discuss in the next section. These statistics are designed to be sensitive to the strength of the pairwise epistatic interactions and the proportion of sites that are drawn from the pairwise epistatic RNA model of Nasrallah and Huelsenbeck (2013). However, it is also possible that the statistics can only detect epistasis if there are a sufficient number of epistatically paired sites in a sequence alignment, or that they are sensitive to the alignment length in general. To investigate all of these possibilities, we simulate alignments on a 3-dimensional grid, defined by *n*_i_ (the number of sites drawn from a site-independent GTR model), and *n*_e_ (the number of sites drawn from the pairwise epistatic RNA model), and the value of *d*, which controls the strength of pairwise epistasis. Briefly, *d* accounts for the strength of epistasis by controlling the relative rate of doublet substitutions, or simultaneous substitutions at both paired (RNA stem) sites, to single substitutions affecting only one paired site. As selection against single mutations (which would break RNA pairing) increases, more double substitutions are expected, and this is captured by a larger *d*. We simulate *d* ∈ {0.0, 0.5, 2.0, 8.0, 1000.0}, which encompasses both realistic and extreme values, and *n*_i_, *n*_e_ ∈ {0, 16, 32, … , 400} (a step size of 16 resulting in a 26 × 26 grid for each value of *d*), excluding the cell in which *n*_i_ = *n*_e_ = 0. This results in alignments that vary in length (16 to 800 nucleotides), in percentage of sites that are epistatically linked (0% to 100%), and in expected percentage of substitutions that are double substitutions (0% to 98.5%). For more information on the simulation procedure, the model, and interpretation of the parameter *d*, please see methods subsections *Model* and *Simulating parameters*. Collectively, we simulate 3375 alignments, which we refer to as our observed alignments.

We consider three posterior predictive test statistics that may detect epistasis: the G93 statistic of Goldman (1993), the maximum of all pairwise (sitewise) mutual information (MI) values, MI_max_, and the kurtosis of these MI values, MI_kurt_. The G93 statistic considers the likelihood of the alignment if sites were drawn identically and independently from a multinomial distribution on all 4^*n*^ site patterns, and has been used as a general diagnostic for model misspecification. Our MI-based measures quantify how similar two site patterns are by comparing the joint distribution on paired states to the product of the sitewise marginal distributions. These measures are sensitive to the co-occurrence of pairs of nucleotides, so for example the 4-taxon alignment (Figure 4E) column ATAT would have a higher mutual information with the column GCGC (where every A co-occurs with a G and every C with a T) than it would GTTG (where there is less frequent co-occurrence). All of these measures are ignorant of the underlying phylogeny, so in practice one must account for its effect via simulation. All three statistics display some sensitivity to *d* on the observed alignments (Figure 1). This means in principle they are all capable of detecting epistasis in alignments.

**Figure 1:**
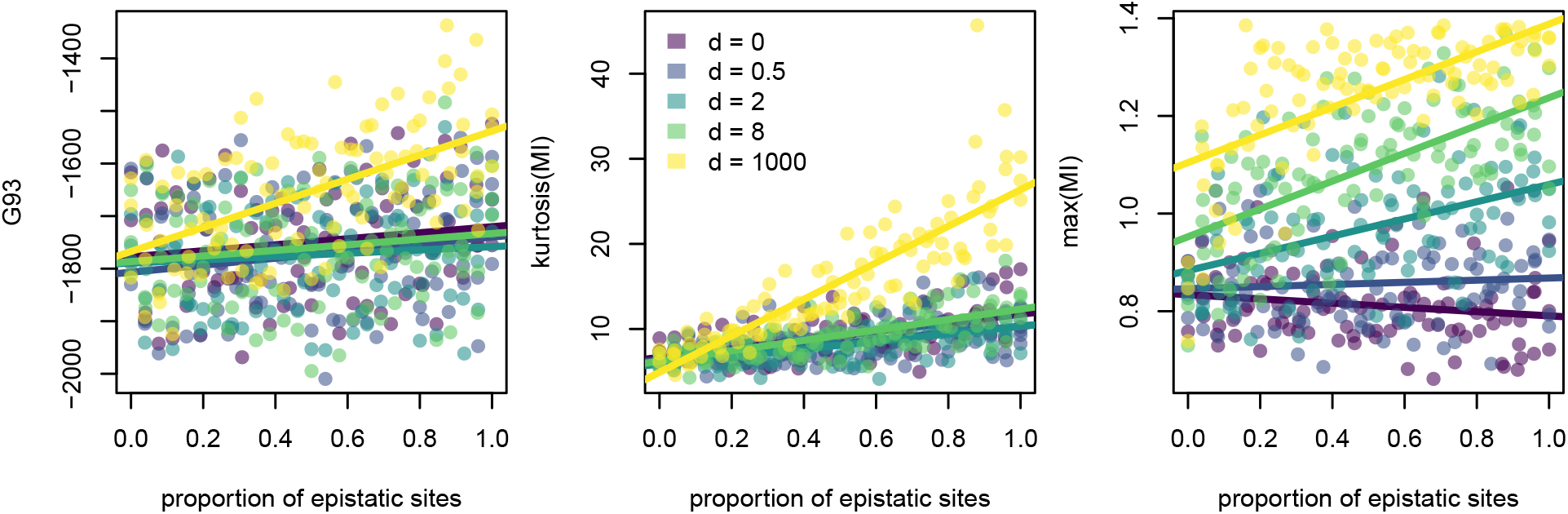
The values of all three test statistics plotted against the proportion of sites in the alignment drawn from the epistatic model, colored by *d*. Values are for the grid of observed alignments corresponding to part (b) of Figure 4. All statistics show at least a degree of sensitivity to the proportion of epistatic sites, and some sensitivity to *d*. The plot is restricted to simulations where 384 ≤ *n*_i_ + *n*_e_ ≤ 416 to remove the effect of the number of sites on the G93 statistic. The results from the full simulation grid are found in Figure S7.

However, the statistics differ in their sensitivity to *d*. Of the three statistics, MI_max_ shows the most consistent separation by *d*, with a steady increase in average value as *d* increases. In contrast, both MI_kurt_ and G93 show large leaps in average values between *d* = 8 and *d* = 1000. Similarly, the MI_max_ shows a consistent relationship of power to the proportion of epistatic sites (*n*_e_/*n*) across all *d* while the G93 and MI_kurt_ statistics which show clear differences between *d* = 8 and *d* = 1000 The G93 statistic is by far the most sensitive to the total number of sites in the alignment, with a correlation coefficient of −0.99 between G93 and *n*_i_ + *n*_e_ where the correlation coefficients for MI_kurt_ and MI_max_ were −0.19 and 0.28 respectively.

### Posterior predictive tests can capture model misspecification due to pairwise interactions between sites

In a posterior predictive framework (Gelman et al., 2004; Brown and Thomson, 2018), the value of a chosen test statistic calculated for the observed alignment is compared to the posterior-predictive distribution of this statistic using posterior predictive p-values. In phylogenetic models which lack closed-form solutions for the posterior predictive distribution, this is accomplished numerically by taking a number of posterior samples, drawing new alignments given the parameter values of each sample, and computing the test statistic on each replicate alignment. The posterior-predictive p-value is then the proportion of replicate test statistics below the observed value. Generally we are interested in two-tailed tests: if the observed value of the test statistic is either extremely large or extremely small, this indicates that the model is not adequately capturing some aspect of the data. We use the test statistics described in the section above to see if a site-iid model is adequate over varying strengths and proportions of epistatic interactions. We use the Bayesian phylogenetic inference software RevBayes (Höhna et al., 2016) to draw samples from the posterior predictive distributions for each of our 3375 observed alignments, allowing us to estimate the posterior predictive p-values for all of our test statistics for each of our observed alignments.

We find that both information theoretic summary statistics outperform G93, with MI_max_ performing the best. An ideal test statistic has a high power (true positive rate) and a false positive rate equal no larger than the specified *α*, which we take to be 0.05. We examine the relationship between the power of each statistic and *d* both by examining power as a function of *d* and the proportion of epistatic sites (Figure 2), and by averaging the power over all proportions of epistatic sites. In both approaches, we find that MI_max_ is the most sensitive to the presence of paired sites in the alignment. The true positive rates in the extreme case of epistatic strength (*d* = 1000) for the G93 (0.18) and the MI_kurt_ (0.56) statistics are matched by the MI_max_ statistic at much lower strengths of epistatic interaction (0.17 at *d* = 0.5 and 0.55 at *d* = 2). Looking over all values of *d* and all observed proportions of epistatic sites, the maximum power of the G93 statistic is only 0.51 while the two information theoretic statistics achieve maximum power of 0.92 and 1.0 for MI_kurt_ and MI_max_ respectively (Figure 2). The MI_max_ statistic has a false positive rate of 0.06, while the MI_kurt_ statistic has a false positive rate of 0.0026, and the G93 statistic has a false positive rate of 0. Overall, the MI_max_ statistic has by far the highest power and the false positive rate does not greatly exceed *α*, making it the best performing statistic we examine. We assess the sensitivity of these results to the strictness of MCMC convergence diagnostics, and find that there is little difference between the results presented here, and those based on either strict or no convergence filtering (Figure S2).

**Figure 2:**
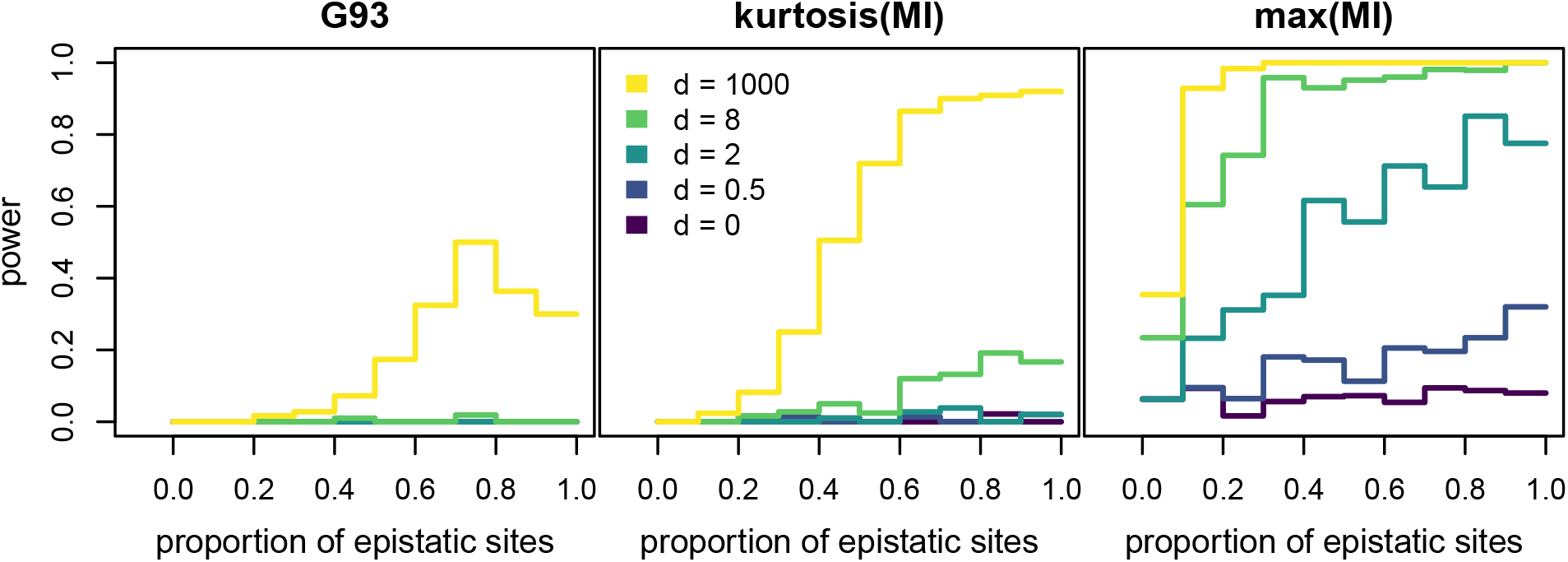
Power to detect epistasis using posterior predictive checks at *α* = 0.05. Power is the proportion tests yielding a statistically significant result, that is, the true positive rate. Curves are averages over windows of 10% of the proportion of epistatic sites.

### Epistatic sites should be retained for phylogenetic inference

For our second question, we want to quantify the effect of our model misspecification by calculating the effective worth of an epistatic site in the context of a site-iid model. Nasrallah et al. (2011) define the effective sequence length, *n*_eff_ to be the length of a hypothetical alignment drawn from a site-independent model which yields the same phylogenetic accuracy as the epistatic alignment. We expand and reframe this analysis to ask the *worth* of an epistatic site in units of independent sites, and define worth in terms of either accuracy or precision. The relative worth, *r*, is a conversion factor that expresses how many independent sites an epistatic site is worth, and thus also allows us to distinguish between different model misspecification scenarios. In the best case, 0 ≤ *r* ≤ 1, and retaining epistatic sites in an alignment improves inference. In the catastrophic case, *r* < 0, and retaining epistatic sites in an alignment leads to worse inference. In either case, it is most likely that the relative worth of a site is dependent on the strength of epistasis, so the relative worth should in fact be a function, *r*(*d*), and not a constant, *r*.

To infer the effective sequence length, we employ semiparametric regression. In this setup, we have 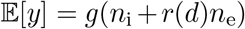, where *y* is a statistic summarizing either inference accuracy or precision and the function *g*() is a third degree I-spline with 5 knots. Third degree (cubic) splines are standard, and model fit is generally robust to the choice of degree and knots (Figure S6). Broadly speaking, accuracy refers to how close estimates are to the true value, and precision is inversely related to how much uncertainty surrounds these estimates (high precision means low uncertainty). The use of semiparametric models like I-splines allows us to avoid specifying a functional form for the relationship between accuracy (or precision) and the alignment size, while still allowing us to compare between datasets with similar accuracies (or precisions) to infer *r*(*d*). For our summary measures, we focus on one accuracy-based measure, the average posterior Robins-Fould (RF) distance to the true tree, and one precision-based measure, the percent of resolved splits in the majority-rule consensus (MRC) tree. In order to quantify the uncertainty in our estimates of *r*(*d*), we use nonparametric bootstrap (Efron, 1992) and fit the model to 100 bootstrap replicate datasets for each summary measure and each value of *d*.

In all cases, we infer that 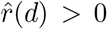, meaning that the model misspecification here falls into the best-case scenario rather than the catastrophic one (Figure 3). For both our accuracy- and precision-based estimates, we infer 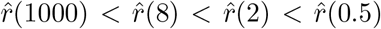, meaning that increasing the strength of epistatic interactions decreases the worth of a site. Additionally, we infer *r*(0) < 1. In the model of Nasrallah and Huelsenbeck (2013), iid sites have a set of stationary base frequencies, *π*_*A*_, *π*_*C*_, *π*_*G*_, *π*_*T*_, while the (paired) epistatic sites have their own set of stationary doublet frequencies, *π*_(*A,A*)_, *π*_(*A,C*)_, …, *π*_(*T,T*)_. So, even without any doublet substitutions, there is model misspecification when assuming a single set of base frequencies, explaining the reduced accuracy at *d* = 0. Oddly, for our accuracy-based modeling, we find that 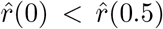. Perhaps most noteworthy though is the fact that, for all values of *d*, we estimate a lower relative worth 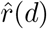 using accuracy than using precision. This means that inference is somewhat more precise than it should be, because the increase in accuracy from adding an epistatic site is smaller than the increase in precision. To give a concrete example, for *d* = 0.5, the relative worth inferred using accuracy is 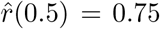, while the relative worth inferred using precision 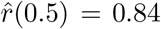. If one added 50 pairs of dependent sites (100 sites total) to an alignment, the accuracy would increase as if 75 independent sites had been added, but precision would increase as if 84 independent sites had been added, a discrepancy of 9 sites.

**Figure 3:**
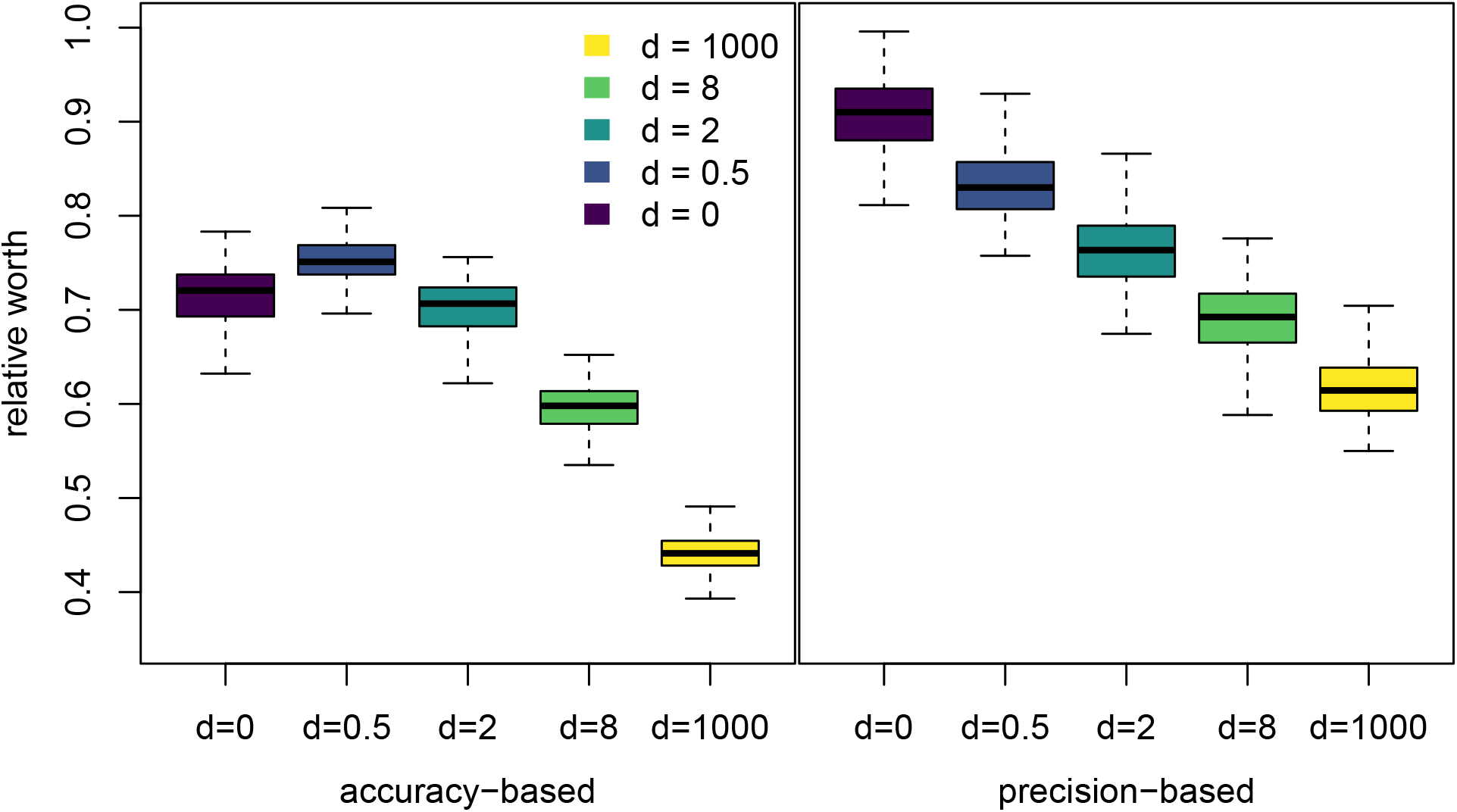
The relative worth of a site, *r*(*d*) estimated using both accuracy and precision. In the best-case model misspecification scenario, 0 ≤ *r*(*d*) ≤ 1, and epistatically paired sites still contribute to phylogenetic accuracy and precision. In the catastrophic model misspecification scenario, *r*(*d*) < 0, and epistatically paired sites decrease phylogenetic accuracy and precision. Inference was performed with semiparametric regression using least squares. Boxplots summarize 100 bootstrap replicates for each value of *d*. Our accuracy measure is the average posterior Robinson-Foulds (RF) distance to the true tree, while our precision measure is the proportion of resolved splits in the majority-rule consensus (MRC) tree. The raw accuracy and precision measure values from the full simulation grid are found in Figure S7.

**Figure 4:**
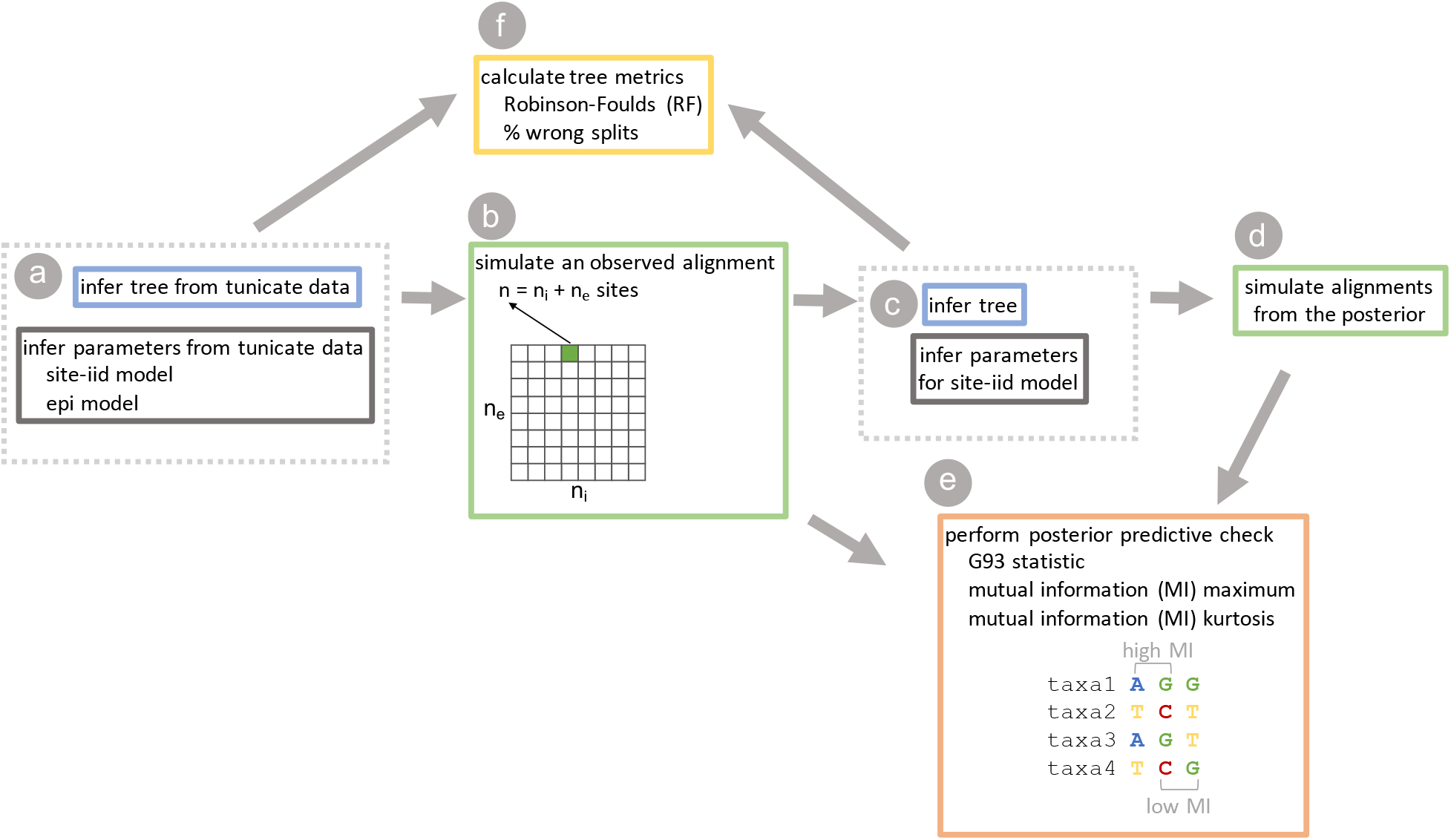
The workflow of our paper. Blue and grey boxes indicate steps with phylogenetic inference, green boxes indicate steps with simulation of alignments. a) We infer a maximum likelihood phylogeny (tree) for the tunicate dataset, and use that to infer parameters for independent sites (site-iid model) and epistatically paired sites (epi model) using the epistatic doublet model of Nasrallah and Huelsenbeck. b) We then simulate 675 alignments of varying numbers of independent (*n*_i_) and epistatic (*n*_e_) sites for each of 5 values of *d* (*d* ∈ {0.0, 0.5, 2.0, 8.0, 1000.0}). We call these the observed alignments. c) For each observed alignment, we use RevBayes to draw samples from the posterior distribution on tree topologies (inferred tree) and GTR model parameters (inferred site-iid model). d) We use RevBayes to sample alignments from the posterior predictive distribution. e) We assess whether any of our three test statistics (G93, MI_max_, MI_kurt_), can detect pairwise epistatic interactions in our observed alignments in a posterior predictive model check. f) We compare the posterior distribution on trees (and summary trees) to the inferred tree to quantify how unmodeled pairwise epistasis affects the accuracy and precision of phylogenetic inference.

We also assessed the sensitivity of our inference of the worth of a site to the MCMC convergence standards employed, and to the choice of summary measure. We find that the overall patterns remain qualitatively unchanged regardless of convergence cutoff (Figure S3). Our alternative accuracy and precision measures broadly concur with the results we have focused on thus far (Figures S4 and S5). All accuracy measures based on the posterior distribution of RF distances to the true tree provide very similar results. The inferred worth is notably lower when defining accuracy based on the proportion of incorrect splits in the MRC tree (though in all cases still positive), and relative worth appears to peak at *d* = 2 rather than *d* = 0.5. Thus, while the posterior distribution shifts closer to the truth with increasing *n*_e_, some incorrect splits retain high support for longer than when increasing *n*_i_. Our alternative precision measure has slightly higher inferred *r*(*d*) than our main measure (15% higher on average), and a number of bootstrap replicates showing 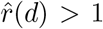. However, we believe this is more likely a sign that the alternative measure (the width of the posterior distribution of distances to the true tree) is for some reason a questionable choice than it is reflecting some actual increase in precision compared to independent sites.

## Discussion

We set out to understand the effect of unmodeled epistasis on phylogenetic inference. Focusing on pairwise epistasis in the form of RNA doublet models, we asked three questions. Can we detect pairwise epistasis in alignments with posterior predictive checks? What is the effect of pairwise epistasis on the quality of phylogenetic inference? If pairwise epistasis is problematic for inference, can we detect it when it is distorting phylogenetic inference? Overall, we find that the right test statistic can detect epistasis with good power (and an appropriate false positive rate), that pairwise epistasis is only mildly harmful to phylogenetic inference, and that it can be detected regardless of how much influence the epistasis has on inference.

### Posterior predictive checks

Our posterior predictive checks for epistasis show that pairwise epistasis can be detected with the appropriate summary statistics. While the standard multinomial likelihood statistic has some ability to detect epistasis at the extreme of *d* = 1000 (an average power across all simulations of 0.17), it cannot detect realistic strengths of epistasis, and so it is not a particularly useful statistic. We also introduced two statistics based on mutual information. In both cases, we compute all the mutual information for all site pairs in the alignment, our statistics simply differ in how they summarize this distribution. One summary we considered, MI_kurt_, is the kurtosis of this distribution, which has decent power at *d* = 1000 (power of 0.54), but at *d* = 8 power is much lower (0.066) and it has little ability to detect weaker epistasis. Our second summary MI_max_, the maximum of the pairwise mutual information values, was much more successful. Averaging over all simulations, power is 0.16 at *d* = 0.5, 0.53 at *d* = 2, 0.87 at *d* = 8, and 0.95 at *d* = 1000, and despite this power the false positive rate is not exaggerated (*α* = 0.05, false positive rate = 0.06, though the distribution of p-values does not appear to be quite uniform under the null hypothesis).

Further research will be needed to understand the specificity of mutual-information-based test statistics and to extend them. While we have shown that the MI_max_ statistic can capture pairwise epistatic interactions, it may also be sensitive to higher-order epistatic interactions, to the presence of context-sensitive mutation, or other dependencies between sites. It is possible that one could identify interacting pairs of sites using MI_max_, and then test if these pairs are significantly interacting using the posterior predictive distribution of mutual information for that pair of sites. Summary statistics that take better advantage over the extreme upper end of the distribution of pairwise mutual information could have even better power to detect pairwise interactions.

### The worth of a site

We defined two scenarios for the worth of an epistatic site. In the catastrophic scenario, epistatic sites are worth some negative number of sites, such that adding epistatically interacting sites to an alignment of independent sites will make inference worse. In the best-case scenario, epistatic sites contribute positively to inference, but not as much as independent sites. Our simulations demonstrate that the sort of epistasis modeled by Nasrallah and Huelsenbeck (2013) falls into the best-case scenario. When epistasis is simulated near maximum strength, we estimate that epistatic sites are worth 45% of independent sites (in terms of accuracy), while more realistic strengths lead to worths of 60% to 75%. One slight caveat is that epistasis has slightly different effects on the accuracy and precision of inference. When we define the worth of a site in terms of the increase in precision, we get slightly higher estimates of worth (62% at the extreme, 70% to 84% for more reasonable values). This means that in practice when epistasis is present in phylogenetic datasets, inference will be slightly overconfident (relative to a dataset without epistatic interactions but with the same accuracy). It also highlights the the fact that high precision does not imply accuracy— strong support for a topology is not a guarantee that the inferred topology is correct. Still, this effect is relatively mild for realistic values of the strength of epistasis, and it does not undermine the simple fact that adding epistatically paired sites improves inference.

### Base frequencies

In our simulations, the model we used for inference was misspecified due to both differences in the relative exchange rates and in the stationary frequencies. We believe this difference may account for the increase in relative worth from *d* = 0 to *d* = 0.5. Our inference model is a standard GTR+G model which works on individual sites, while our simulated data includes a mix of sites drawn from a site-iid GTR model and an epistatic doublet model. The doublet model is inherently a model of site pairs, both because it uses doublet stationary frequencies and because *d* allows for substitutions at both sites. However, the doublet model still produces a pattern of substitutions at individual sites, so we can decompose the model misspecification into two parts: the portion due to paired substitutions, and the portion due to the fact that there are essentially two different site-level GTR models. The misspecification due to paired substitutions is a strictly increasing function of *d*: the higher *d* is, the larger the proportion of all substitutions that are doublet substitutions, and this is completely unaccounted for by the site-independent model. The misspecification due to the difference in underlying GTR models (the real one underling the independent sites, and the hypothetical one underlying the epistatic sites) is somewhat less transparent. However, it appears that the difference is large between the two GTR models at *d* = 0, decreases until *d* = 2, then increases with *d*. The interaction of these two forms of misspecification could help explain our observation that the worth of an epistatic site at *d* = 0 is less than for *d* = 0.5, but similar to *d* = 2.

### Alternative summaries and other studies

Empirical applications of epistatic inference models have shown that epistatic and site-independent models infer different trees from the same dataset. Meyer et al. (2019) compared inferences using their model of pairwise dependence to inference with GTR+G on a number of real datasets, and found that the inferred trees were quite different. One possible interpretation of this is that there is bias introduced by using models of independent sites on epistatic datasets. However, our simulations suggest that another possible explanation for this phenomenon is simply that to site-iid models the effective sequence length is smaller than the real sequence length, and thus estimates from site-iid will vary more about the truth than estimates made with epistatic models. If this is the case, then there are two valid approaches for obtaining better estimates of a phylogeny in the presence of epistasis: building better models and/or using more data Philippe et al. (2011). Both approaches have drawbacks and are not entirely independent. Better models will be more difficult and computationally expensive to fit, may not be as general-purpose, and the increase in parameters may require an increase in data to fit. Adding more data means that issues of heterogeneity of the evolutionary process across sites and loci become more pronounced, which may require more parameter-rich models even if epistasis is itself unmodeled.

Previous simulations have suggested that unmodeled epistasis can be relatively problematic for inference. Nasrallah et al. (2011) used simulations and found that accuracy was reduced by as much 50% in the presence of epistasis, and calculated effective sequence lengths that were frequently 17%-33% of the true length. Our results suggest that the choice of summary measure for defining the worth of a site is key to understanding this discrepancy. For example, our precision-based measures both produce more generous estimates of the relative worth of an epistatic site than our accuracy-based measures. Perhaps more importantly, though, our accuracy results change somewhat if we change our definition of accuracy to be based on the posterior or on a single summary tree. Using accuracy measures based on the posterior distribution of RF distances to the true tree, we estimate 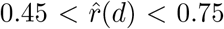 for all values of *d* considered, and 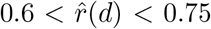 for more realistic values. But using the MRC tree alone to define accuracy (based on the proportion of splits in the MRC not in the true tree), we instead infer 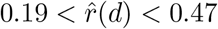 for all values of *d*, and 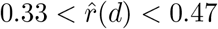 for more realistic values. These values are more closely aligned to the results of Nasrallah et al. (2011), who focused on point estimates of the phylogeny (specifically the maximum likelihood phylogeny).

Thus it seems that while epistatic sites are relatively helpful for shifting the posterior distribution of tree closer to the truth, they are less useful for removing incorrect splits from the MRC tree. The issue of excess precision is exacerbated when focusing in on the summary tree: the relative worth based on MRC tree precision is about twice as high as the relative worth based on MRC tree accuracy. To give a concrete example, at *d* = 0.5, using precision we estimate 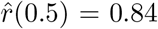, while using our alternative accuracy measure we estimate 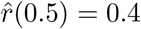. Thus, an alignment of 500 iid and 500 independent sites would have an MRC tree with the same resolution as an entirely iid alignment of an alignment of 920 sites, but that tree would have the same number of incorrect splits as an alignment of 700 sites. We thus urge caution in interpreting MRC trees from real datasets, given the presumed prevalence of epistasis. Further, we suggest that where possible, inferences about evolution integrate out the phylogeny, as the overall posterior distribution on phylogenies is much less strongly affected by the presence of unmodeled pairwise epistasis.

### Practical applications

As dependencies between sites are likely common in real data, our work has practical significance to phylogeneticists. Model checking via posterior predictive simulations (or the parametric bootstrap for maximum likelihood inference) is always advisable. We suggest that when checking for model violations, researchers include the MI_max_ statistic. The detection of dependencies between sites does not necessarily invalidate inference. Rather, if it reveals the presence of dependent evolution, researchers need to be aware that their effective sequence length is smaller than the real sequence length. Consequentially, the estimated phylogeny may be farther from the truth than expected while uncertainty around that phylogeny is likely underestimated, and results should be interpreted accordingly.

### Other forms of model misspecification

In this paper, we assume that every site evolves along one single phylogeny of interest, and that the model misspecification is due to unmodeled pairwise epistatic interaction. Many other forms of model misspecification exist, which can make phylogenetic inference difficult (see for example, Philippe et al., 2011). Long branch attraction can lead inference astray even with otherwise correct substitution models, and forces such as recombination, horizontal gene transfer, and incomplete lineage sorting mean that real sequence alignments may have multiple underlying phylogenies. It is possible that the notion of relative worth could be useful in comparing the severity of these, and other issues. For some problems, the relative worth will be positive, and more sequence data will improve phylogenetic accuracy. In other cases, the relative worth may become negative, in which case more sites will not solve the problem. In general, more strongly negative relative worth causes more difficulty for phylogenetic inference.

### Conclusion

Overall, our results are quite promising for phylogenetic inference in the face of unmodeled pairwise epistasis. While pairwise epistasis decreases the accuracy and precision of inference, it does not do so catastrophically. The addition of epistatic site pairs to an alignment will still lead to overall better inference, simply not inference as good as adding an equal number of independent sites from the true model. Moreover, when there is pairwise epistasis in an alignment we have shown that it can reliably be detected with a new test statistic: the maximum pairwise mutual information of sites. Thus, if one is worried that pairwise epistasis is interfering with their estimates, they can now detect it with good power and a low risk of false positives. Further work will need to be done using other models of epistasis (pairwise and higher order) to check that our findings on the effect of epistasis are not simply localized to one region of the space of epistatic models. Additional investigations examining the effect of the tree length could also prove useful. It is likely that our new test statistic will be useful for detecting other forms of pairwise epistasis, though this should also be tested. Our results suggest that in practice, phylogenies inferred from alignments with pairwise epistasis are still reliable and valid estimates.

## Methods

### Model

For our simulations, we employ the epistatic RNA doublet model of Nasrallah and Huelsenbeck (2013), which we implemented in RevBayes (Höhna et al., 2016). In this model, there are two categories of sites: paired sites evolving dependently (which we will refer to as epistatic sites) and unpaired sites evolving independently (which we will refer to as independent sites). Instead of directly parameterizing a selective coefficient against single mutations, the model parameterizes an enrichment factor for doublet substitutions which controls how common doublet substitutions are relative to single substitutions. Within each category, it is assumed that all sites (or site pairs) evolve under the same model. The independent sites evolve under a standard GTR+G substitution model (Tavaré, 1986). The epistatic sites evolve under an extension of the GTR+G model to the state space of doublets. We abbreviate these models as iid and epi, respectively. The (symmetric) nucleotide exchange rates and the shape parameter of the gamma-distributed rate heterogeneity are shared between epi and iid models. We now detail the epistatic model.

For any pair of coupled sites, let ***Q***^e^ denote the instantaneous rate matrix describing changes from doublet ***x*** = (*x*_1_, *x*_2_) to doublet ***y*** = (*y*_1_, *y*_2_), where 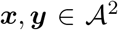 and 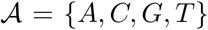 is the character set. Without loss of generality, let the first element refer to the more 5’ nucleotide of the pair. The rate matrix has elements

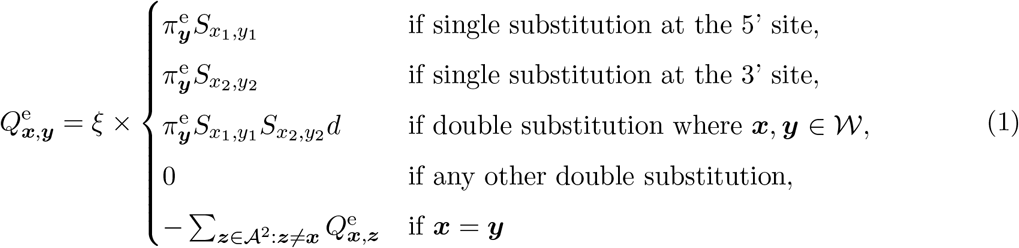

where ***S*** is the GTR exchangeability matrix (shared with the independent sites) with *e.g.* 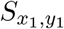 the rate of exchangeability between nucleotides *x*_1_, *y*_2_ ∈ {*A, C, G, T*} (by definition, ***S*** is symmetric), 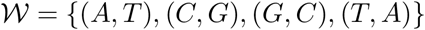 is the set of Watson-Crick pairs, ***π***^e^ = (*π*_(*A,A*)_, *π*_(*A,C*)_, …, *π*_(*T,T*)_) are the stationary state frequencies of the 16 possible doublet states, *d* is the relative rate of double to single mutations between doublets, and *ξ* is a rate-scaling factor that normalizes ***Q***^e^ to one substitution per single site for comparability with the independent sites. As with mutation-selection models on codons (*e.g.* Yang and Nielsen, 2008), this approach localizes dependencies between sites by working on groups of sites assumed to be mutually independent. However, in the epistatic doublet model, simultaneous substitutions at multiple sites are allowed, and selection is modeled only implicitly, through the relative frequency of doublet substitutions, *d*.

For completeness, we now review the rest of the model. For any non-coupled sites, let ***Q***^i^ denote the instantaneous rate matrix describing changes from state *x* to state *y*, where 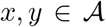 is the character set of nucleotides as above. The rate matrix has elements

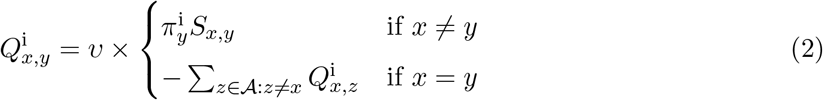

where ***S*** is the GTR exchangeability matrix as above, shared with the epistatically paired sites, ***π***^e^ = (*π*_*A*_, *π*_*C*_, *π*_*G*_, *π*_*T*_) are the stationary state frequencies of the 4 nucleotides, and *υ* is a rate-scaling factor that normalizes ***Q***^i^ to one substitution per site.

The likelihood of the multiple sequence alignment is computed under the standard phylogenetic assumption that a pair of epistatic sites is independent of all other pairs of sites (and all unpaired sites), and that an unpaired site is independent of all sites. Thus, the only site-to-site dependencies are those modeled in ***Q***^e^, and we can write the likelihood of the alignment as,

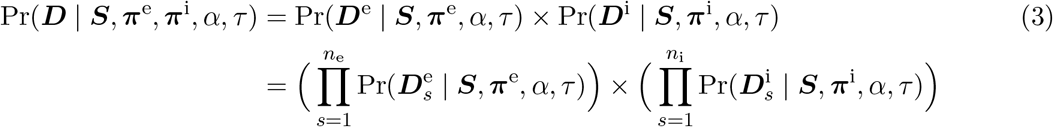

Where ***D***^e^ and ***D***^e^ are the subalignments consisting of all epistatically paired sites and all independent sites, respectively, *α* is the shape (and rate) parameter of the Gamma-distributed among-site rate variation, *τ* is the tree topology with branch lengths, and the per-site-pair and per-site like-lihoods, 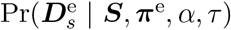 and 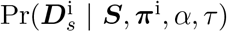, are computed using the Felsenstein pruning algorithm (Felsenstein, 1981) with the rate matrices ***Q***^e^ and ***Q***^i^ as defined above.

The parameter *d* controls the strength of epistatic interactions. At *d* = 0, there are no double substitutions, and if both *d* = 0 and 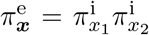, the epistatic model collapses to the standard site-independent GTR+G model. Taking the limit *d* → ∞ and *ξ* → 0 such that the expected number of substitutions is constant, all substitutions (at paired sites) are double substitutions. With 0 < *d* < ∞, a portion of substitutions at epistatically paired sites are expected to be doublet substitutions while the remainder will be single-site changes. We can compute the expected fraction of the substitutions that are doublet substitutions as

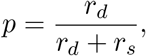

with *r*_*d*_ defined as

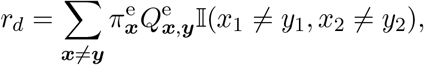

(where 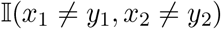 is an indicator for doublet substitutions) and *r*_*s*_ defined as

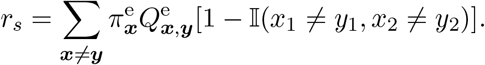

Both *r*_*d*_ and *r*_*s*_ depend on *d* through the normalization constant *ξ*, a dependency which we can remove by multiplying *p* by *ξ*^−1^/*ξ*^−1^, giving us

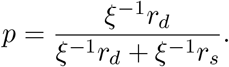

From the definition of ***Q***^e^, we can see that, holding ***S*** constant, *ξ*^−1^*r*_*d*_ increases linearly in *d* while *ξ*^−1^*r*_*s*_ is a constant in *d*. It can thus be seen that, for fixed values of other substitution parameters, the proportion of doublet mutations is a sigmoid function of log *d*.

### Simulation

#### Parameter grid

With the epistatic doublet model, there are two variables that govern the capacity for epistasis to affect phylogenetic inference: the strength of the interactions, and the proportion of the alignment that is epistatically paired. To investigate the effect of the strength of epistasis, we simulate with *d* ∈ {0.0, 0.5, 2.0, 8.0, 1000.0}, which encompasses both realistic (see below) and extreme values. These values correspond to 0%, 11.2%, 33.6%, 66.9%, and 99.6% of all substitutions being doublet substitutions at a pair of sites. The scaling constant *ξ* in ***Q***^*e*^ ensures that the expected number of substitutions per site remains constant across all values of *d*. To understand the effect of adding epistatic sites, for each value of *d* we simulate a grid where we independently vary the number of epistatic sites, *n*_e_ and independent sites, *n*_i_. This setup is more informative than one where we hold the total number of sites constant while varying the proportion of paired sites because it allows us to examine the sensitivity of posterior predictive tests to both proportion and absolute number of paired sites. By comparing the accuracy of tree inference before and after adding epistatic sites, we can assess if we are in the catastrophic model misspecification regime (inference is degraded by the addition of epistatic sites) or the best-case regime (inference is improved by adding epistatic sites). For each value of *d*, we simulate an alignment from a fixed tree at each cell of a parameter grid defined by *n*_i_, *n*_e_ ∈ {0, 16, 32, …, 400}, excluding the cell in which *n*_i_ = *n*_e_ = 0. At each grid point we simulate a single alignment, and we refer to these alignments collectively as the *observed* alignments. We note that *n*_e_ must be even because epistatic sites are simulated as (nonoverlapping) coupled site-pairs.

#### Simulating parameters

To ensure that our simulation regime is realistic, we target our simulating parameters on values inferred from the tunicate dataset of Tsagkogeorga et al. (2009). This dataset is an 18s rRNA alignment for 110 taxa, including 95 species of tunicates. We downsample (randomly) to 50 ingroup species and infer a tree in RAxML (Stamatakis, 2014) using a single GTR+G model. Fixing the tree to the RAxML tree, we use RevBayes (Höhna et al., 2016) to infer the parameters of the epistatic doublet model. For the RevBayes analysis, we split the alignment into a subalignment containing only unpaired (loop) sites (682 sites total) and a subalignment containing only paired (stem) sites (644 total nucleotides in 322 site-pairs). The paired sites are recoded such that AA,AC,…,TT are treated as RevBayes-style standard characters 0,1,…,F. For simulations, we use this RAxML tree and the posterior mean parameter values from the RevBayes analysis. These parameters include the underlying GTR model exchange rates, the underlying GTR model stationary frequencies, the stationary frequencies of the doublets, and the shape parameter of the gamma-distributed rate heterogeneity. All continuous parameter values are available in Supplementary Table S1, and Figure S1 depicts the tree used for simulation. The tree length is 4.58, meaning that on average there should be approximately 4.58 substitutions at each column in the simulated alignments.

Our simulating values of *d* aim to include both extreme values and biologically relevant values. We infer a value of *d* of 0.65 for this tunicate dataset. Nasrallah and Huelsenbeck (2013) inferred values of *d* of 7.59 and 9.72 on a dataset spanning eukaryotes for analyses fixing and inferring the tree respectively. We therefore chose simulating values of *d* ∈ {0.5, 2.0, 8.0} to cover the range of values inferred from datasets of natural sequences. Simulating *d* = 0 allows us to disentangle the effect of model misspecification due to the doublet stationary frequencies from model misspecification due to paired substitutions. Finally, simulating *d* = 1000 allows us to consider the extreme regime where almost all (98.5%) substitutions at paired sites are doublet substitutions.

#### Bayesian inference

We use RevBayes (Höhna et al., 2016) to infer unrooted phylogenies for each of the observed alignments. In all cases, we use a single GTR+G substitution model, intentionally ignoring the presence of epistasis in the datasets. Details of the model setup, priors, and MCMC moves used are available in the supplement. For each analysis we run two independent MCMC chains. We exclude runs that fail convergence diagnostic tests from downstream analysis to avoid analysis artifacts. As we are interested in the phylogeny specifically, our convergence diagnostics focus on the tree and branch lengths. First, we use the average standard deviation of split frequencies (ASDSF) to compare topologies between the two chains. To account for the possibility of branch-length convergence issues, we also check whether the tree length distributions differ between chains by using the potential scale reduction factor (PSRF, Brooks and Gelman, 1998). However, we must balance the stringency of our convergence standards against the number of analyses that must be discarded. To this end, we discard all chains where either ASDSF > 0.05 or PSRF > 1.1, and in the supplement we present additional results assessing sensitivity of our results to different convergence standards.

### Posterior predictive assessments

The posterior-predictive distribution is the distribution on new (replicate) datasets, ***D***^rep^, that we could draw from our posterior distribution, ℙ(*θ* | ***D***). It is obtained by integrating the probability of a new dataset given a particular value of the model parameters ℙ(***D***^rep^ | *θ*), across the posterior

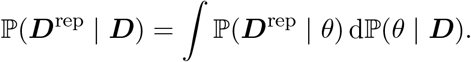

Often we are more interested in a particular feature of a dataset, given by a test statistic *T*(***D***). The posterior predictive distribution for this test statistic is given by,

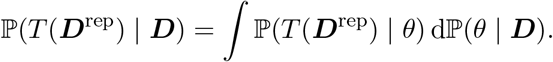

We can use the posterior predictive distribution of a test statistic to determine whether our model is adequate, or in other words whether it fits the data in the absolute sense. If the model is adequate, then the observed value of the test statistic, *T* (***D***), is consistent with the posterior predictive distribution ℙ(*T* (***D***^rep^) | ***D***). We can thus compute a posterior predictive p-value, ℙ(*T* (***D***^rep^) < *T* (***D***) | ***D***), and if the posterior-predictive p-value is smaller than some threshold *α*, we declare the model to be inadequate. In practice, one obtains a Monte Carlo estimate of the posterior predictive p-value by simulating new datasets using the draws from the posterior obtained by MCMC, computing the test statistic for each, and calculating the proportion greater than or equal to the observed value. For a more complete introduction to posterior predictive model checks see Gelman et al. (2004), or for a review of model adequacy in evolutionary biology see Brown and Thomson (2018).

One of our key questions is, can the presence of unmodeled epistatic interactions be detected with posterior predictive checks? A test statistic should be chosen on the basis of its ability to detect a particular form of model misspecification. As epistasis has yet to be studied from a posterior predictive perspective, there are currently no posterior predictive approaches designed explicitly to detect it. We first examine whether a standard phylogenetic posterior predictive test statistic—the multinomial likelihood test statistic of Goldman (1993)—can detect this model misspecification. We then develop two new information theoretic statistics that directly address the expected behavior of epistasis.

We examine the performance of these test statistics on simulated datasets in two ways. First, by using only the simulated observed alignments, we examine if these test statistics are sensitive to the proportion of epistatic sites in an alignment or to *d*. Next, we perform posterior predictive checks for all inferred phylogenies to determine if the statistics are able to detect epistasis in practice. As an example of this distinction, consider that in principle the GY statistic is capable of detecting the difference between certain GTR (Tavaré, 1986) models and certain HKY (Hasegawa et al., 1985) models. However, if we simulate an alignment under HKY and infer it under GTR, we will not see any evidence of misspecification (Bollback, 2002), so in practice it cannot detect all forms of misspecification.

#### Multinomial likelihood

The multinomial likelihood test statistic treats the multiple sequence alignment ***D*** as a draw from a multinomial distribution on all possible site patterns. For *t* taxa and a DNA or RNA alignment, there are 4^*t*^ such patterns, and for an alignment of length *n* we observe some number *s* ≤ *n* site patterns. If *s*_*i*_ is the number of times we see site pattern *i*, the maximum likelihood estimate of the multinomial probability of seeing pattern *i* is *p*(*i*) = *s*_*i*_/*n*. The test statistic is simply the log-likelihood of this dataset using these estimated probabilities, or 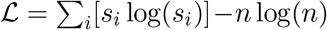.

#### Mutual information

Under a phylogenetic model, all sites are conditionally independent given the phylogeny. However, under the epistatic doublet model, epistatic pairs of sites evolve dependently along the phylogeny due to both *d* and the use of doublet stationary frequencies. In particular, the presence doublet substitutions make it more likely that when one site in a pair changes along a branch, the other site does as well. This should result in pairs of epistatic sites having more similar patterns than independent sites, and the degree of this similarity should depend on *d*. What is needed, then, is a way to capture this idea of the similarity of sites.

Mutual information (MI) is a measure of the dependency of two variables, quantifying the amount of information that one contains about the other. The higher the MI, the more that knowing the value of one variable tells you about the other, in other words the more similar the two variables are. As above, we assume that we have an alignment ***D*** consisting of *t* taxa and *n* sites, and here we again denote the alphabet of this alignment 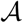. The MI of a pair of sites indexed by (*i, j*) is given by,

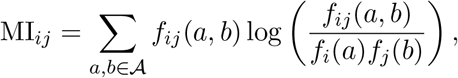

where *f*_*i*_(*a*) is the relative frequency of character *a* at site *i*,

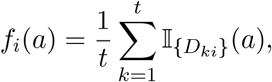

and *f*_*ij*_(*a, b*) is the relative joint frequency of character *a* at site *i* and character *b* at site *j*,

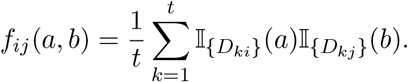

In non-phylogenetic contexts, MI has been used to predict RNA secondary structure from multiple sequence alignments (see Freyhult et al., 2005, and references therein).

As MI is ignorant of the phylogeny, we need additional context to interpret its value. If we had an *a priori* hypothesis about a pair of interacting sites (*i, j*), we could compute the posterior predictive distribution of MI_*ij*_ and compare the observed value to this distribution. The posterior predictive distribution accounts for similarity due to shared evolutionary history, so if the observed value is larger than the posterior predictive distribution, there must be some other factor at play, such as pairwise epistasis. However, if one wants to test for the presence or absence of pairwise epistasis at the level of the entire alignment, this will not work. Instead, we can compute the MI of all pairs of sites. As we expect epistasis to increase the MI between pairs of interacting sites, the upper tail of the distribution of pairwise MI values should be informative with respect to the overall presence and strength of epistasis. We consider two summaries of this distribution. First we consider its kurtosis, which should be sensitive to the strength of epistasis and proportion of epistatic sites, as the more interactions the more values that should fall in the right tail and the stronger the interactions the larger those values should get. We also consider the max, which should be sensitive to the presence or absence of epistasis, but is less likely to be sensitive to the proportion of epistatic sites.

#### Computing the relative worth of an epistatic site

Let *y* be a statistic summarizing the accuracy or precision of inference on a dataset, and (*y* | *n*) denote conditioning on the number *n* of sites in the alignment. The sites could be drawn from a site-independent model, or a model that introduces dependence among sites, such as the Nasrallah-Huelsenbeck model of pairwise epistasis. As examples *y* could include the distance between the maximum likelihood tree and the true tree, or the variance of the posterior distribution of trees. The effective sequence length is *n*_eff_ such that 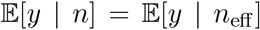. That is, if we repeatedly draw datasets with *n* sites from some arbitrary, possibly pairwise dependent, model and *n*_eff_ sites from the (site-independent) model used for analysis, then *n*_eff_ is the number of independent sites such that the average accuracy or precision is the same as for the *n* sites from the arbitrary, possibly misspecified, model. If *y* is a statistic that summarizes the accuracy of inference, then this definition includes the effective sequence length of Nasrallah et al. (2011). Note that we can ignore the number of taxa because it must be the same between the datasets. It is also worth noting that this formulation of an effective sequence length is broader than the formulation of the effective sample size. Classically, the notion of effective sample size, *n*\* is defined with respect to the variance of an estimator 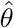. Specifically, the effective sample size is *n*\* such that 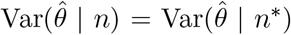. The variance of an estimator is given by 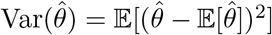. Thus, if we take 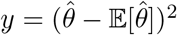, then we can see that our definition includes the standard definition of effective sample size.

In our case, with *n* = *n*_i_ + *n*_e_, we can define a model for the effective sequence length as *n*_eff_ = *n*_i_ + *r*(*d*)*n*_e_ where *r*(*d*) is the relative worth of an epistatic site compared to an independent one for a given value of *d*. The dependency on *d* is necessary because we expect that as *d* increases and more substitutions are paired, the relative worth should decrease. It is possible that, for a given value of *d*, *r*(*d*) is also not a constant; if there is asymptotic bias and the estimated tree does not converge to the true tree, then *r*(*d*) must eventually go to 0. However, in our finite data regime there is no evidence of asymptotic biases, and a model with constant *r*(*d*) will suffice. Estimating *r*(*d*) is complicated by the fact that we do not know an appropriate functional form for 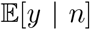 for any of our metrics. We know that accuracy and precision should both increase with increasing *n*_eff_, but how rapidly this happens is unknown. Thus, to infer *r*(*d*) we turn to models where we do not need to explicitly specify this relationship. All modeling is done in R (R Core Team, 2018). Specifically, we use I-splines (Ramsay, 1988) to model the relationship between *y* and *n*_eff_ as 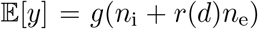 where *g* is a spline. We use the R package splines2 to obtain the basis matrix for the splines, using degree 3 splines with 5 knots. Preliminary analyses suggest that the inferred relative worth is generally stable if the degree and/or number of knots are changed. To ensure that *g*() is monotonically increasing, the coefficients that are fit to this basis matrix must be non-negative. We use log and logit transformations to keep our response variables unbounded, taking negatives where necessary to make them monotonically increasing rather than monotonically decreasing. For this purpose, we employ the R package penalized (Goeman, 2010), which allows us to estimate maximum likelihood (least squares) coefficients with bounds on the coefficients. By estimating *r*(*d*), we can determine whether we are in the catastrophic model misspecification regime (*r*(*d*) < 0) or the best-case regime (0 ≤ *r*(*d*) ≤ 1) for any value of *d*.

#### Accuracy

As we are interested in Bayesian inference, we are not primarily interested in the accuracy in point estimates of the phylogeny but in the overall goodness of the posterior distribution of trees. As our distance measure, we employ the Robinson-Foulds (RF) distance (Robinson and Foulds, 1981). RF distance is a purely topological measure between a pair of trees, capturing the number of splits (bipartitions of taxa) present in one tree but not in the other. The quantity that we are interested in is thus the posterior distribution on distances to the true tree. Given that we have samples from our posterior distribution on phylogenies, we can obtain samples from the posterior distribution on tree distances. We consider multiple univariate summaries of this distribution, namely the mean, median, minimum, and maximum of the distances.

While comparisons based on the entire posterior distribution are useful for understanding the overall performance of inference, they do not necessarily reflect the experience of practitioners inferring phylogenies. In practice, the posterior distribution is, at least for the purposes of visualization, generally reduced to a single summary tree, often a majority rule consensus (MRC) tree. Thus, as an alternative accuracy measure, we take the proportion of splits in the MRC tree that are not in the true tree as an error measure of the point tree estimate.

#### Precision

To investigate precision-based effective sequence lengths, we must define a measure of the precision or variance of our posterior distributions. In the best-case scenario, where epistatic sites are simply less informative than independent sites, one would intuitively expect that the variance of the posterior distribution on trees should increase. On the other hand, in the catastrophic scenario it is possible that there is an increase in information about certain edges in the tree, and the variance of the posterior distribution may actually decrease. While the variance of a phylogeny is defined (*e.g.* Willis, 2019), the time required to compute this variance makes it prohibitively expensive for our purposes (Brown and Owen, 2019). In order to address the matter of variance, we thus turn to two surrogates.

The first metric we consider is the resolution of the majority-rule consensus (MRC) tree. The MRC tree is obtained by including all splits in the posterior that occur with a frequency above 50%. As the amount of information in an alignment increases, there should be more splits with sufficient signal to place in the MRC tree and the MRC tree should include more splits. An MRC tree on *t* taxa includes a maximum of 2*t* − 3 non-trivial splits, so dividing the number of (non-trivial) splits in the MRC tree by 2*t* − 3 produces a standardized value in [0,1] which we call the proportion of resolved splits. At 0, the MRC tree is completely unresolved (a star tree), while at 1 it is a fully resolved tree. While this metric takes a somewhat circuitous path to precision, the focus on the summary tree ties it more closely to tangible effects of variance encountered when reading a paper that estimates a phylogeny.

As an alternate metric, we consider the width of the 95% CI of the distances to the true tree. The more information there is in an alignment, the narrower we expect the posterior distribution on trees, and thus the narrower we expect the distribution on distances to the true tree. We focus here on the RF distance as it is a purely topological measure and does not include any potentially confounding effects due to erroneously long (or short) estimated trees. This metric has the downside of being linked to the accuracy of the inference, which is less than ideal, but like our other metric it is correlated with the true precision of the distribution, and it has the benefit of requiring no extra computations, somewhat reducing the otherwise heavy CO_2_ cost of this paper.

## Data and code availability

Code and data to reproduce this study’s results can be found at https://github.com/wsdewitt/phyload.

## Acknowledgments

We thank our PhD advisors (Frederick Matsen IV, Vladimir Minin, Jesse Bloom, and Kelley Harris) for the freedom to pursue this project. We also thank Frederick Matsen, Vladimir Minin, and Joseph Felsenstein for their many helpful comments on earlier versions of this manuscript. We thank two anonymous reviewers for feedback that improved the manuscript. W.S.D. thanks Mike Harms for helpful discussions.

A.F.M. was supported by National Science Foundation grant DGE-1762114 and the ARCS Foundation Fellowship. W.S.D. was supported by the National Institutes of Health via a NRSA fellowship from the National Institute of Allergy and Infectious Diseases (F31AI150163), and National Institute of General Medical Sciences (R01 GM113246, PI Matsen).

