## Supplemental Text for "Robustness of phylogenetic inference to model misspecification caused by pairwise epistasis"

Supporting Information

### Simulating parameters

**Table S1:** Simulating parameters for the alignments.

| Parameter | Value | Role |
|---|---|---|
| $\pi_i$ | [0.33, 0.21, 0.24, 0.22] | Stationary frequencies for the iid sites (in alphabetical order) |
| $\pi_e$ | [0.02, 0.02, 0.02, 0.16, 0.02, 0.02, 0.19, 0.01, 0.01, 0.25, 0.02, 0.05, 0.12, 0.01, 0.06, 0.02] | Stationary frequencies for the epistatically-coupled site pairs (in alphabetical order) |
| $\alpha$ | 0.35 | Shape and rate parameter for Gamma-distributed among-site rate variation |
| $s$ | [0.10, 0.20, 0.10, 0.11, 0.36, 0.13] | Relative exchange rates between nucleotides (the lower triangular portion of $\boldsymbol{S}$) |

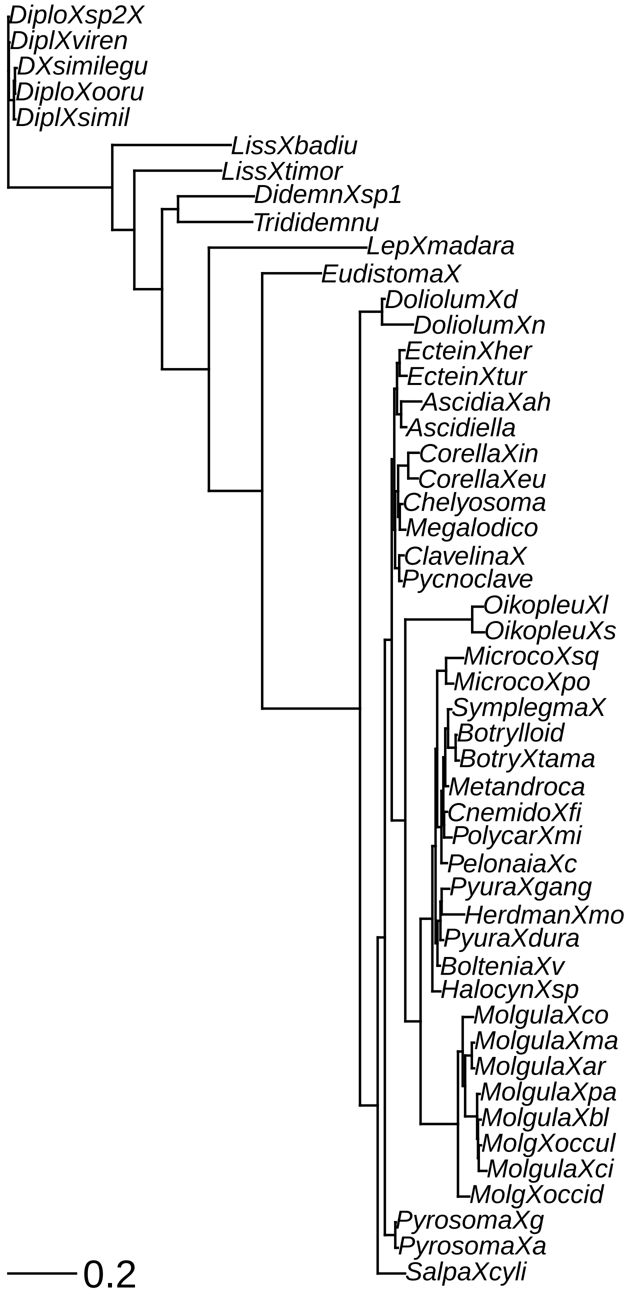


**Figure S1:** The tree inferred in RAxML used for both estimating the epistatic doublet model parameters in RevBayes and for simulation. Scale bar is 0.2 substitutions per site.

### Sensitivity to MCMC convergence

All analyses presented in the paper filter out simulations for which MCMC convergence failed. We define failure to be PSRF $>$ 1.1 or ASDSF $>$ 0.05, a threshold that reflects our desire to balance convergence standards against retaining sufficient simulation replicates from which to make inferences. To determine whether our results are robust to the convergence criterion, we now present results for Figures 2 and 3 in which we consider using all the data regardless of MCMC convergence diagnostics ("no convergence"), the standards presented in the main text ("convergence") and a stricter standard (PSRF $>$ 1.01 or ASDSF $>$ 0.01, "strict convergence"). As can be seen in Figure S2, the inferred power to detect epistasis is unaffected by convergence standards. Similarly, Figure S3, the inferred worth of epistatic sites is qualitatively unaffected by convergence cutoffs. The inferred worth values are also similar with the exception of the precision-based worth using strict standards. For $d = 1000$, the worth inferred from precision is higher than for the other convergence standards. However, the stricter convergence standards resulted in discarding over 33% of the simulations at $d = 1000$, while for the other $d$ values less than 25% were discarded, so the discrepency could simply be a result of losing data.

### Bayesian analyses of simulated data

All analyses of simulated data are done in RevBayes. We employ a GTR+G substitution model, with flat Dirichlet priors on the symmetric exchange rates and base frequencies, and a half-Cauchy(0,1) prior on the alpha parameter for gamma-distributed among-site rate variation. The tree prior is uniform over topologies. For flexibility, we use a hierarchical branch length prior which parameterizes the branch lengths in terms of a prior on the expected tree length. Specifically, we define $\theta$ to be the expected tree length, and let $\theta \sim \text{Lognormal}(0.0, 1.17481)$ distribution, such that $\theta$ has a prior median of 1, with a 95% prior CI of [1/10,10]. We define $\lambda := n_{\text{branches}}/\theta$, and all branch lengths are iid Exponential($\lambda$). We run the analysis for 20000 iterations after a burnin of 2000 iterations, using two independent chains. Our move scheme uses on average 139 moves per iteration, with 127 moves on the tree (5 SPR moves, 25 NNI moves, and 97 branch length scaling moves) and 12 moves on continuous parameters. This is equivalent to 3,058,000 MCMC generations in programs like MrBayes or BEAST, discarding the first 9.1% as burnin.

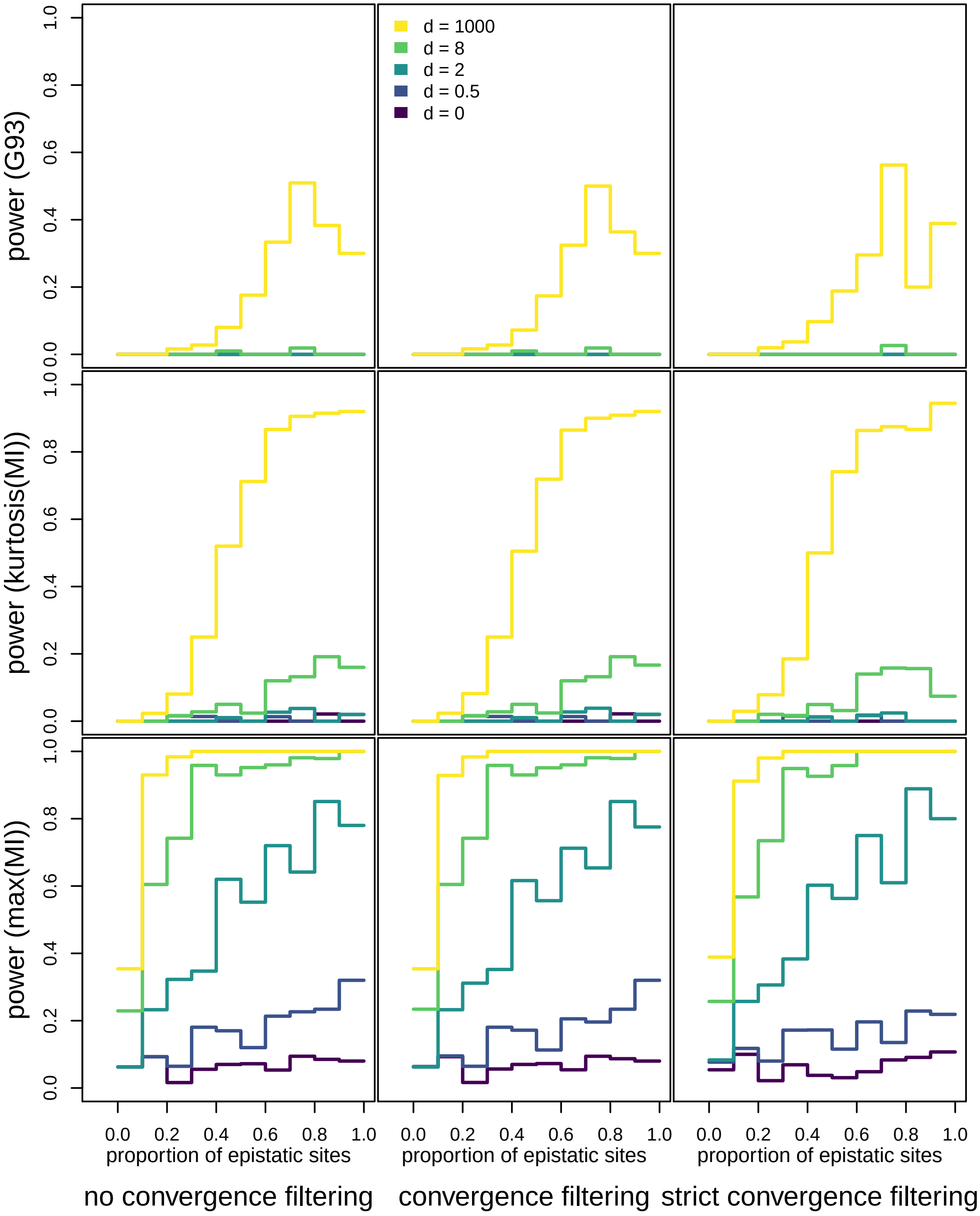


**Figure S2:** Power to detect epistasis using posterior predictive checks at $\alpha = 0.05$ across three different convergence thresholds. The lowest threshold is to simply include all simulations ("no convergence"), the intermediate threshold is to remove any runs with PSRF $> 1.1$ or ASDSF $> 0.05$ ("convergence"), and the strict threshold is to remove any runs with PSRF $> 1.01$ or ASDSF $> 0.01$ ("strict convergence"). Power is the proportion tests yielding a statistically significant result, that is, the true positive rate. Curves are averages over windows of 10% of the proportion of epistatic sites.

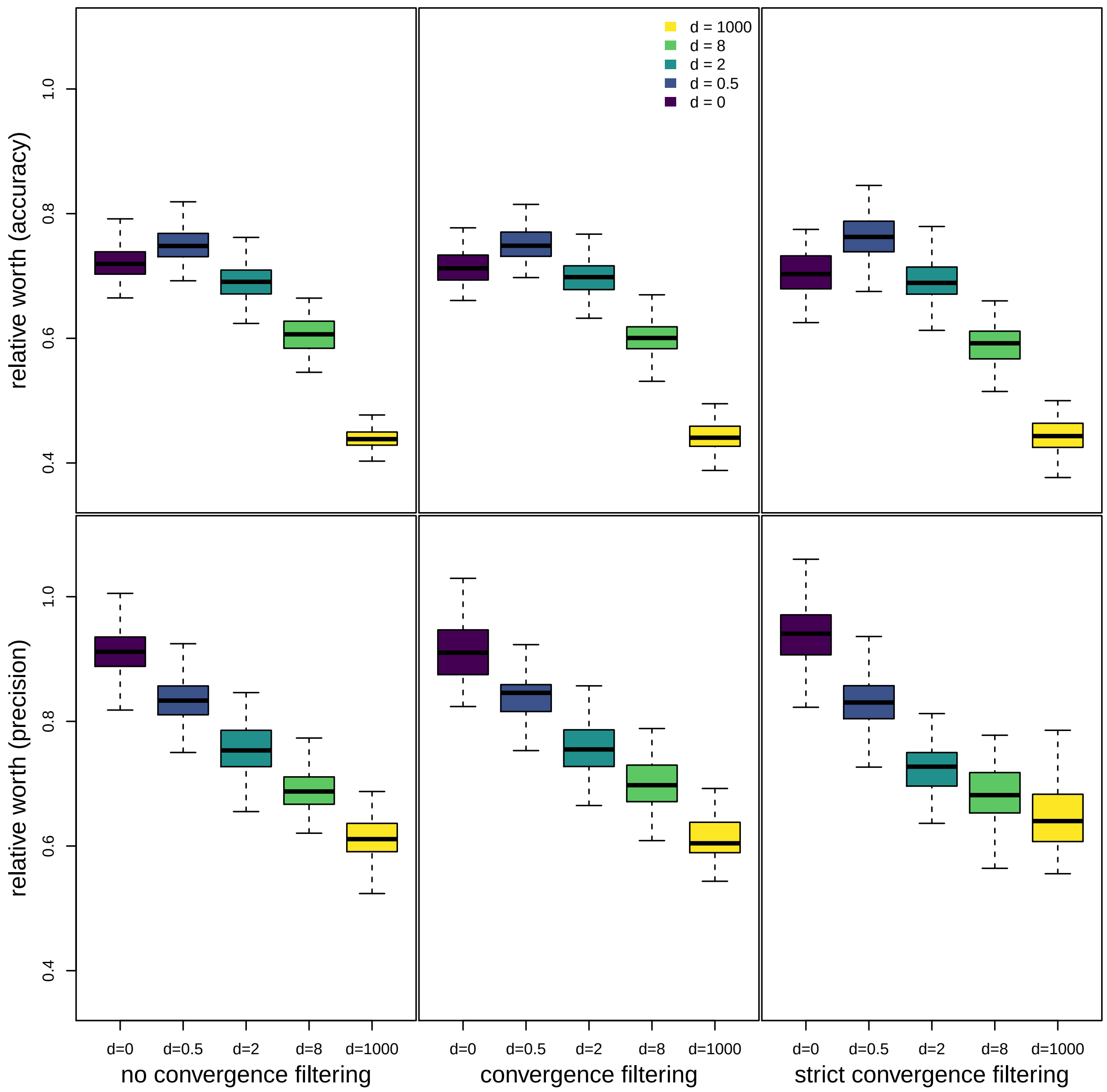


**Figure S3:** Bootstrapped estimates of $r(d)$ for our simulated values of $d$ across three different convergence diagnostic thresholds. The lowest threshold is to simply include all simulations ("no convergence"), the intermediate threshold is to remove any runs with PSRF $> 1.1$ or ASDSF $> 0.05$ ("convergence"), and the strict threshold is to remove any runs with PSRF $> 1.01$ or ASDSF $> 0.01$ ("strict convergence"). Inference of worth was performed via least squares using semiparametric regression. Accuracy-based estimates use the average posterior RF distance to the true tree, while precision-based estimates use the proportion of resolved splits in the MRC tree. The boxplots summarize 100 bootstrap replicates.

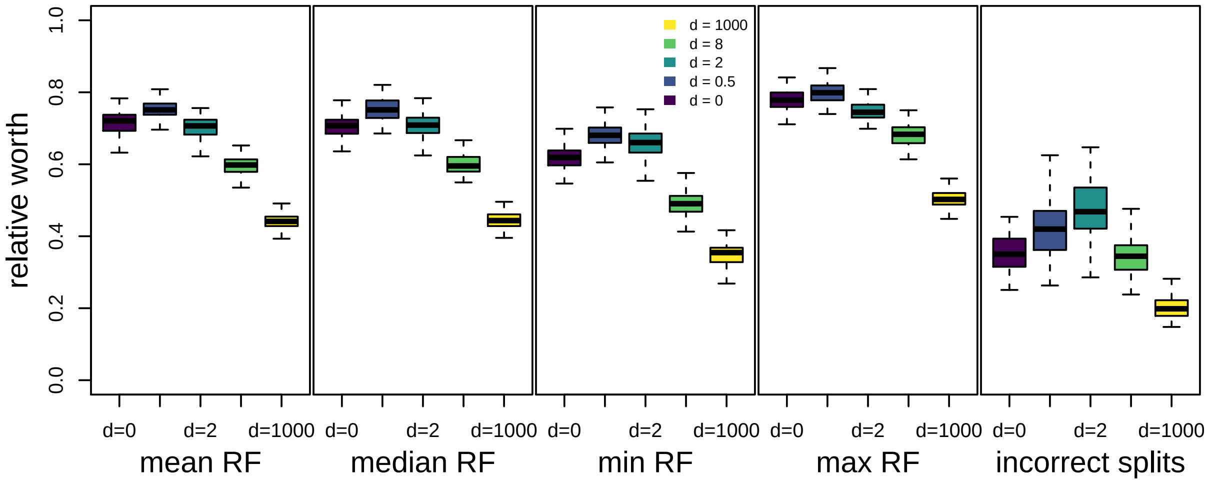


**Figure S4:** Bootstrapped estimates of $r(d)$ for our simulated values of $d$ for several measures of accuracy. The left four panels are different summaries of the posterior distribution of RF distances to the true tree, in order the mean, median, minimum, and maximum. The right-most panel measures accuracy by (one minus) the proportion of resolved splits in the MRC tree that are not in the true tree. Inference of worth was performed via least squares using semiparametric regression. The boxplots summarize 100 bootstrap replicates.

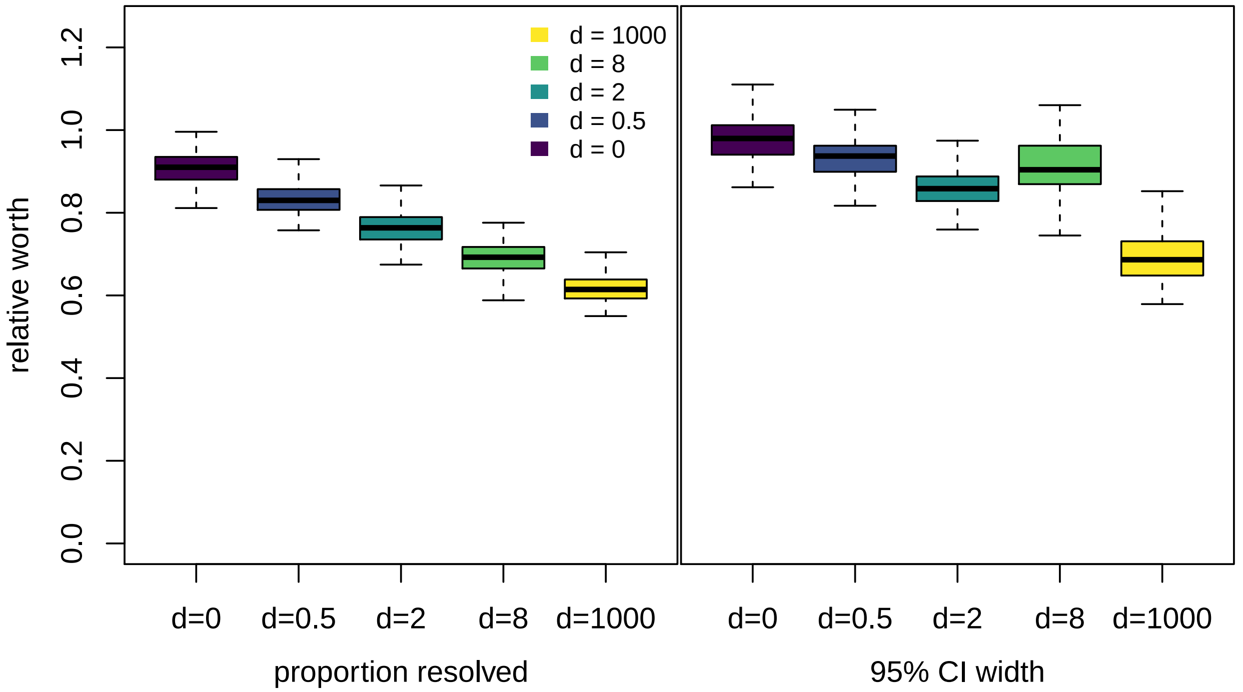


**Figure S5:** Bootstrapped estimates of $r(d)$ for our simulated values of $d$ for two different measures of precision. On the left, precision is measured using the proportion of resolved splits in the MRC tree. On the right, precision is measured as the width of the 95% CI of the posterior distribution of RF distances to the true tree. Inference of worth was performed via least squares using semiparametric regression. The boxplots summarize 100 bootstrap replicates.

### Spline model

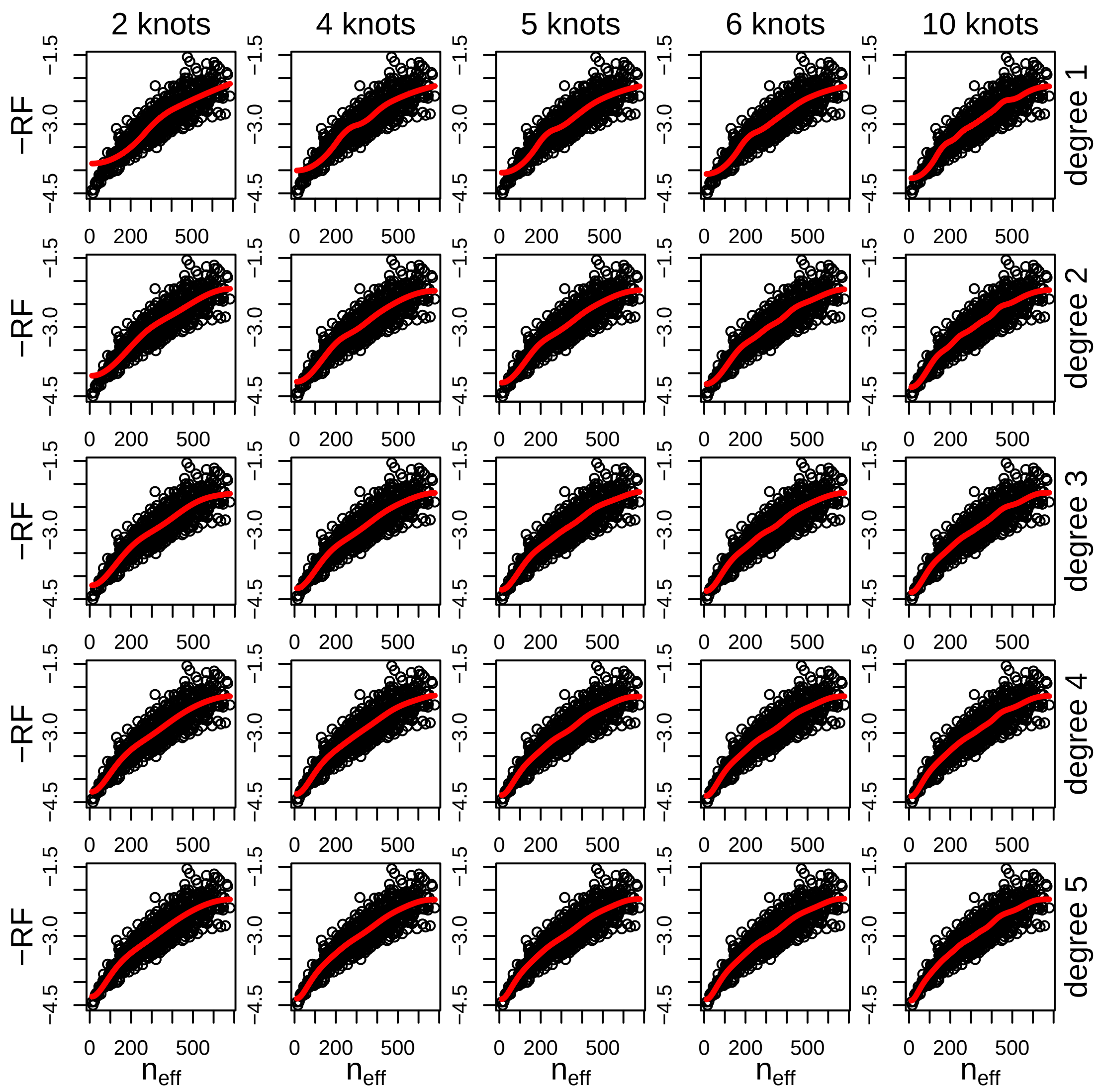


**Figure S6:** Fit of Ispline models for a variety of numbers of knots and spline degree for $d = 2$. The x-axis is the effective sequence length, $n_{\text{eff}}$, which is a function of the value of $r(d)$ inferred for each plot. In our analyses, we use 5 knots and 3rd degree polynomials, which is the center plot. The best fit line is qualitatively similar for analyses with models that use similar numbers of parameters as our chosen model. The x-axis is relatively stable across all plots, indicating similar inferred values of $r(d)$.

### Test statistics and tree metrics across the full grid

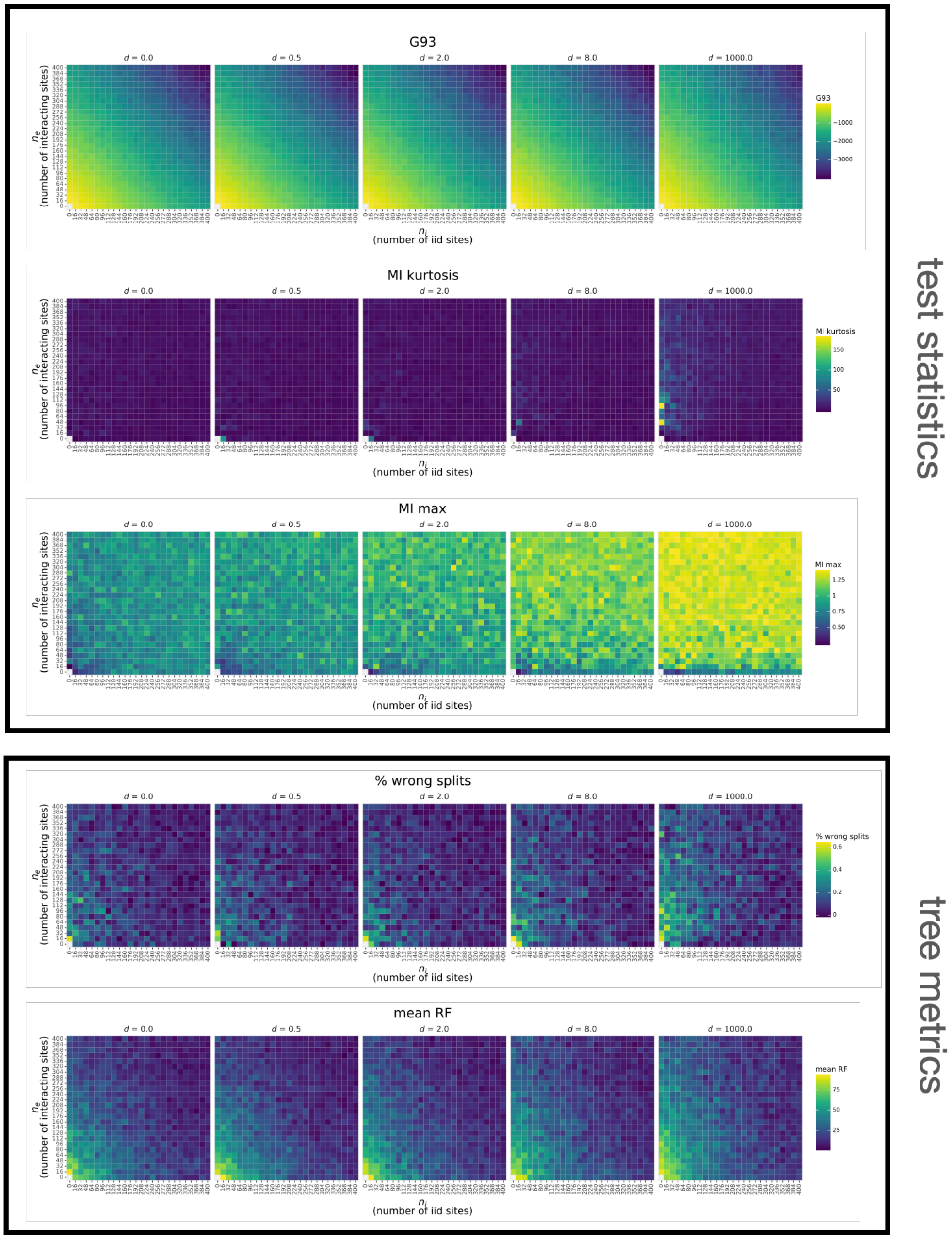


**Figure S7:** Heatmaps showing test statistic and tree metric values for the full simuation grid. Top panels show the three alignment-based test statistics used in the posterior predictive check (Figure 4E): the statistic from Goldman, 1993 (G93), the maximum mutual information value (MI max) and the kurtosis of the mutual information values (MI kurtosis). A portion of the values in these grids (specifically the main and adjacent off-diagonals) are shown as a function of the proportion of sites that are epistatic in Figure 1. Bottom panels show the two metrics used to compare (Figure 4F) the tree inferred by the site-iid model (Figure 4A) to the tree used to simulate the alignemnts (Figure 4C): proportion of splits in the MRC tree that are not in the true tree (% wrong splits) and the mean Robinson-Foulds distance from posterior tree samples to the true tree (RF mean). These values are used to estimate relative worth as shown in Figure 3. Note the color scale varies from metric to metric.
